# Correlating viscosity and molecular crowding with fluorescent nanobeads and molecular probes: *in vitro* and *in vivo*

**DOI:** 10.1101/2022.06.27.497768

**Authors:** Sarah Lecinski, Jack W. Shepherd, Kate Bunting, Lara Dresser, Steven D. Quinn, Chris MacDonald, Mark C. Leake

## Abstract

In eukaryotes, intracellular physicochemical properties like macromolecular crowding and cytoplasmic viscoelasticity influence key processes such as metabolic activities, molecular diffusion, and protein folding. However, mapping crowding and viscoelasticity in living cells remains challenging. One approach uses passive rheology in which diffusion of exogenous fluorescent particles internalised in cells is tracked and physicochemical properties inferred from derived mean square displacement relations. Recently, the crGE2.3 Förster Resonance Energy Transfer (FRET) biosensor was developed to quantify crowding in cells, though it is unclear how this readout depends on viscoelasticity and the molecular weight of the crowder. Here, we present correlative, multidimensional data to explore diffusion and molecular crowding characteristics of molecular crowding agents using super-resolved fluorescence microscopy and ensemble time-resolved spectroscopy. We firstly characterise *in vitro* and then apply these insights to live cells of budding yeast *Saccharomyces cerevisiae*. It is to our knowledge the first time this has been attempted. We demonstrate that these are usable both *in vitro* and in the case of endogenously expressed sensors in live cells. Finally, we present a method to internalise fluorescent beads as *in situ* viscoelasticity markers in the cytoplasm of live yeast cells, and discuss limitations of this approach including impairment of cellular function.

## 1. Introduction

The cytoplasm of live cells is highly dynamic and undergoes physicochemical regulation throughout the cellular lifespan in response to external stress such as glucose starvation, and internal stress such as ageing (1,2). Mainly composed of water, it contains nutrients, proteins, and ions, which must interact with each other to perform the basic chemical reactions that sustain life. The *physical* state of the cytoplasm is therefore as important as its chemical state – in too viscous a solution key diffusive processes, such as those involved in signalling pathways (3,4), will slow down, and in too crowded an environment protein folding may be compromised despite the presence of chaperone proteins (5,6). The viscoelasticity of the cytoplasmic matrix also performs a mechanical role for the cell as a whole, providing internal pressure and resistance to mechanical deformations (4,7–9).

Studying viscoelasticity in single cells via light microscopy tracking of probes, or through applying forces, is known as *microrheology*. To measure mechanical properties of living cells, various techniques have been used, including assessment by atomic force microscopy (AFM) (10–15) and magnetic twisting cytometry (16,17). Force transduction tools such as magnetic and optical tweezers can either stretch a cell (18,19) or controllably perturb a measurement bead in so-called active microrheology (20,21) to measure the response to high and low frequency deformations. A microplate rheometer may be used for the study of cell cultures such as human tissues (22). Particle tracking microrheology in cells may be used without the need to perturb marker beads (23,24). Here, the diffusion of a particle through the internal cellular landscape is observed through optical microscopy. The resulting particle track, or *trajectory* as it may also be denoted, has a property known as the mean square displacement (MSD), which essentially measures the average spatial separation of the trajectory at time *t* and *t+dt* for all *dt*. This characteristic of the trajectory may then be signal-transformed numerically using the inverse Laplace transform, and then Fourier transformed to give a complex number characteristic of the trajectory (25). The real and imaginary parts of this are known as *G’* and *G”* respectively and relate to the physical properties of conservative elastic trapping in the matrix, and dissipative viscous diffusion - the trajectory and MSD are directly related to the sheer creep compliance, which is a measure of deformability. For example, a sheer creep compliance slope of 1 indicates a perfectly viscous solution. A requirement for this type of analysis is that the absolute physical environment is estimable only from tracking one particle over a sufficient timescale, with sufficient temporal resolution.

Measuring molecular crowding, on the other hand, is significantly more challenging. The crowding environment cannot be estimated from diffusion measurements of typically micron length scale diameter marker beads because the excluded volume effects are on the length scale of a single protein, *i.e*., a few nm in diameter. The experimental state-of-the art is to use a Förster Resonance Energy Transfer (FRET)-based probe to measure crowding. One such that we have utilised previously is called crGE2.3 (26–28), consisting of a donor fluorophore, mGFP and acceptor, mScarlet-I, linked by a flexible amino acid hairpin, forming an efficient FRET pair system as reported and characterised in the literature (29–31). As molecular crowding increases, the donor and fluorophore are pushed closer together, and the FRET signal increases (26,32,33). However, how this FRET readout is correlated to macroscale diffusive behaviour is currently not well characterised or understood.

Particle-tracking microrheology using intracellular fluorescent beads has been used to probe the viscoelastic properties across a wide range of conditions in various cells types, including plant and mammalian cells (34–36). Budding yeast, *Saccharomyces cerevisiae* is an invaluable model for molecular crowding studies, however, no methodology currently exists using this model. Nonetheless, we note that recent development were made reporting viscoelastic properties of the cytoplasm of the fission yeast (*S. pombe*) and its influence on microtubules dynamics (37) a type of yeast cell dividing through symmetrical fusion events as opposed to the asymmetrical cell division of the budding yeast used in this study. Here, we report our efforts to combine the two methodologies *in vitro* and *in vivo* using a range of fluorescent beads and the crGE2.3 crowding sensor expressed in budding yeast. First, we characterise *in vitro* diffusion of both sensors and beads of diameters from a few tens to a few hundreds of nm, with crGE2.3 FRET efficiencies in a range of different molecular weight crowders, in order to construct a standard against which we can compare *in vivo* results. We then report the same quantities *in vivo* for the endogenously expressed sensor. Finally, we introduce a method for transforming fluorescent beads into live yeast.

## 2. Methods

### 2.1 Slimfield microscopy

High spatial and temporal precision Slimfield microscopy (38) was performed on a bespoke fluorescence single-molecule microscope described previously (39,40). For the yellow-green and red bead experiments, single wavelength lasers were used (488 nm wavelength OBIS LS, Coherent, and 561 nm OBIS LS, Coherent, respectively) in epifluorescence illumination mode. Collected fluorescence emission light was imaged on a BSI Prime 95B camera after expansion to 53 nm/pixel at an exposure time of 2.44 ms. The camera region of interest was cropped such that the total frame-to-frame time was approximately 5 ms. In each experiment, we used 1 mW laser power delivered at the back aperture of the objective lens, 16 MHz 12-bit depth mode and high sensitivity gain.

For *in vitro* sensor experiments, we first collected 10 frames using millisecond time scale alternating laser excitation (ALEX), using the same exposure time as for *in vivo* imaging with no significant lag time between consecutive frames, to excite both the donor and the acceptor, followed by 1,000 image frames of single-colour imaging of the acceptor only. As the acceptor does not undergo FRET to the donor, all energy is emitted as visible light and the molecule is therefore maximally visible. Here, both lasers were set to 20 mW incident at the back aperture of the objective lens, but all other imaging settings were the same as for beads. *In vivo* imaging parameters were identical to those of the *in vitro* sensors, unless stated otherwise.

### 2.2 Single particle tracking

Micrographs were analysed using our previously developed PySTACHIO software suite (41). We did not calculate molecular stoichiometries or fluorescence intensities, concentrating only on the diffusive behaviour. For all images we used struct_disk_radius=10 and min_traj_length=3. For bead experiments the signal-to-noise ratio cutoff (snr_filter_cutoff) was set to 0.6, and was set to 0.4 for single sensor experiments.

### 2.3 *In vitro* diffusion assays

#### CrGE2.3 Sensor freely diffusing in tunnel slides

Molecules were imaged *in vitro* during their diffusion in a tunnel slide constructed with two pieces of double-sided tape, a standard microscopy slide, and a 22×22 mm #1.5 coverslip (Menzel Glaser) as described previously (42). 20 μl of a 1/200 dilution in imaging buffer (10 mM NaPi sodium phosphate) at pH 7.4 supplemented with relevant concentration of glycerol or Ficoll 70 or 400 of 5 μm uncoated carboxy-latex beads (Micromer-01-02-503, Micromod) were added to the slide and incubated in an inverted hygrostatic chamber for 5 minutes to allow the beads to non-specifically bind to the coverslip, the presence of these beads allowing to locate the coverslip surface and facilitate focus prior imaging. Unbound beads were then washed out with 200 μl 10mM NaPi (pH 7.4), the imaging buffer. 20 μl BSA (bovine serum albumin) at 1 mg/ml in water was then added to the tunnel and incubated at room temperature for five minutes to passivate the glass surfaces. Remaining BSA was then washed out with 200 μl of imaging buffer (10mM NaPi, pH 7) with or without the concentration of interest in glycerol or Ficoll and 20 μl of fluorescent protein was added (1:20 dilution with appropriate imaging buffer from a stock solution of *ca*. 0.2 μg/ml). Finally, the tunnel slide was sealed with nail varnish to prevent drying of the flow-cell prior to imaging.

#### Purification of the crGE2.3 sensor for *in vitro* analysis

The crGE2.3 sequence holding the mGFP-mScarlet-I FRET-pair was introduced into a PRSET-A expression plasmid by ligation using NdeI and HindIII as restriction enzymes. The introduced sequence holds a 6HIS tag sequence in the open reading frame and is attached to the N-terminal region of the sensor, and incorporating a His-tag specific purification. Expression of the protein was performed following an adapted protein purification protocol (26). Briefly, the plasmid was transformed into BL21(DE3) *Escherichia coli* strain then grown at 37°C to mid-log phase (OD600 between 0.6 and 0.8) in LB medium supplemented with ampicillin at 1 mg/ml. The cells were then induced with 100 μg/ml of Isopropyl β-D-1-thiogalactopyranoside (IPTG) and left to grow for 24h at 18°C. The culture was centrifuged at 4000 rpm for 20 minutes, and resuspended in lysis buffer (10 mM NaPi, 100 mM NaCl, 0.1mM AEBSF (4-(2-aminoethyl)benzenesulfonyl fluoride hydrochloride), pH 7.4), then left to freeze at −80°C, then left to thaw on ice before performing for sonication while still on ice (Sonopuls HD 2070, Bandelin, 100% power burst, effective power of 70 W, 6 cycles of 20 s pulse followed by 20 s pause). The lysate was centrifuged for 40 minutes at 15,000g, the pellet was discarded and lysate supernatant filtered with a 0.45 μm filter. A HIS-tag purification was performed using a commercial gravity nickel column (HIS-Gravitrap, GE-healthcare). An additional size column exclusion purification was performed using a Superdex 200 10/300 GL column (Amersham Biosciences) in 10 mM NaPi, pH 7.4. Expression and purification were analysed by 4-12% SDS-PAGE gel. Concentrations were measured using the absorbance at 280 nm wavelength on a Nanodrop 2000 Spectrometer (ThermoFisher), with 56395 M^-1^cm^-1^ (cited here to 5 significant figures) for the crge2.3 extinction coefficient obtained from the sensor amino-acid sequence via the Expasy-ProtParam tool using the Edelhoch method via the extinction coefficients for Tryptophan and Tyrosine determined by Pace methods (43–47). Finally, the aliquoted protein (at a concentration of approximately 12 μg/ml) was directly flash frozen with liquid nitrogen and stored at −80 °C.

#### Fluorescent carboxy-latex beads freely diffusion in tunnel slides

Tunnel slides were prepared as above. Rather than purified sensor,20 μl of a 1/10,000 dilution of carboxylate latex fluorescent beads (Fluospheres, Invitrogen) of either 20 nm (F-8786), 100 nm (F8803) or 200 nm diameter (F-8810) was introduced. The tunnel slide was then sealed with nail varnish prior to imaging.

### 2.4 *In vivo* imaging sample preparation

Budding yeast cells expressing the crGE2.3 sensor were grown to mid-log phase (OD600 0.4-0.6) overnight at 30°C with shaking in growth synthetic minimal media buffer (a commercial formula following supplier recommendation, composed of YNB, Yeast Nitrogen Base (LDT Formedium, UK), supplemented with synthetic amino-acid drop-out lacking histidine (LDT Formedium, UK), 2% glucose). 1 ml of this culture was then centrifuged at 1000g for five minutes, the supernatant discarded, and the pellet resuspended in buffer (20 nM Tris HCl pH 7.5) prior to loading on tunnel slides for imaging. Tunnel slides were made with tape and plasma cleaned cover slips, and incubated first with 1 mg/ml Concanavalin A (ConA) for 5 minutes. This was washed out with 200 μl phosphate buffered saline (PBS, pH 7), and 20 μl 2 mg/ml BSA was introduced and incubated for five minutes to passivate the remaining glass surfaces. Finally, 20 μl of the resuspended yeast were introduced and incubated for 5 minutes inverted to bind to the ConA, then washed out with 200 μl and the slide was sealed prior to imaging.

### 2.5 Time-correlated Single Photon Counting (TCSPC) Fluorescence Lifetime measurements and analysis

Fluorescence decays of both free GFP and crGE 2.3 were acquired using a FluoTime300 single photon counting spectrophotometer with hybrid photomultiplier tube (PMT) detector (PMYA Hybrid 07, Picoquant). Samples were prepared in a volume of 200 μl 10mM NaPi at pH7.4 containing the crGE2.3 FRET sensor diluted 1/200 into the appropriate concentration of glycerol, Ficoll-70 or Ficoll-400. As shown in Fig. 4A, samples were placed into a quartz cuvette (Starna Scientific Fluorimeter Micro Type 22-Q-3). To obtain time-resolved fluorescence, samples were excited by pulsed excitation at 485nm wavelength with a repetition rate of 50 MHz (LDH-P-C-485, Picoquant). Fluorescence decays were recorded peak emission for GFP of 509 nm at the magic angle and until a peak of 10,000 photon counts were collected. Fluorescence decays were then fitted after reconvolution with the instrument response function using a tri-exponential model (Fig. 4B), where It is the intensity at time, *t*, normalized to the intensity at *t* = 0, and *t_i_* and *a_i_* represent the fluorescence lifetime and fractional amplitude of the *i^th^* decay components, as previously (48).

**Figure 1:**
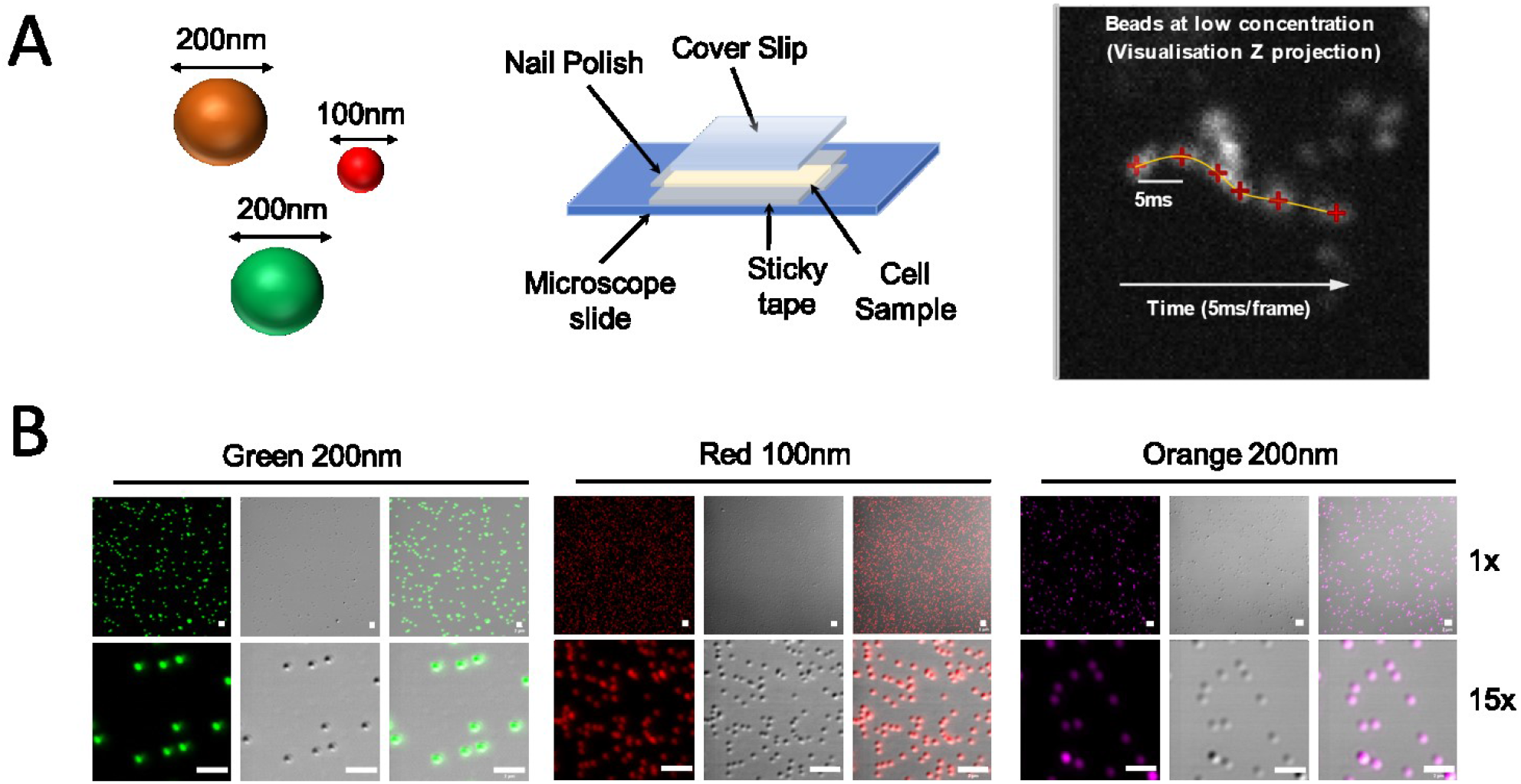
Experimental set up to image fluorescently tagged beads in live budding yeast cells. (A) Schematic showing fluorescently labelled beads of varying sizes and experimental set up for in vitro imaging by confocal and Slimfield microscopy. Left to right: the beads used; construction of a tunnel slide; and an indicative fluorescent trace from a Slimfield micrograph of diffusing beads. (B) Representative confocal images of different fluorescent microspheres using a 63x oil immersion objective lens at 1x and 15x zoom. All scale bars: 2 μm.

**Figure 2:**
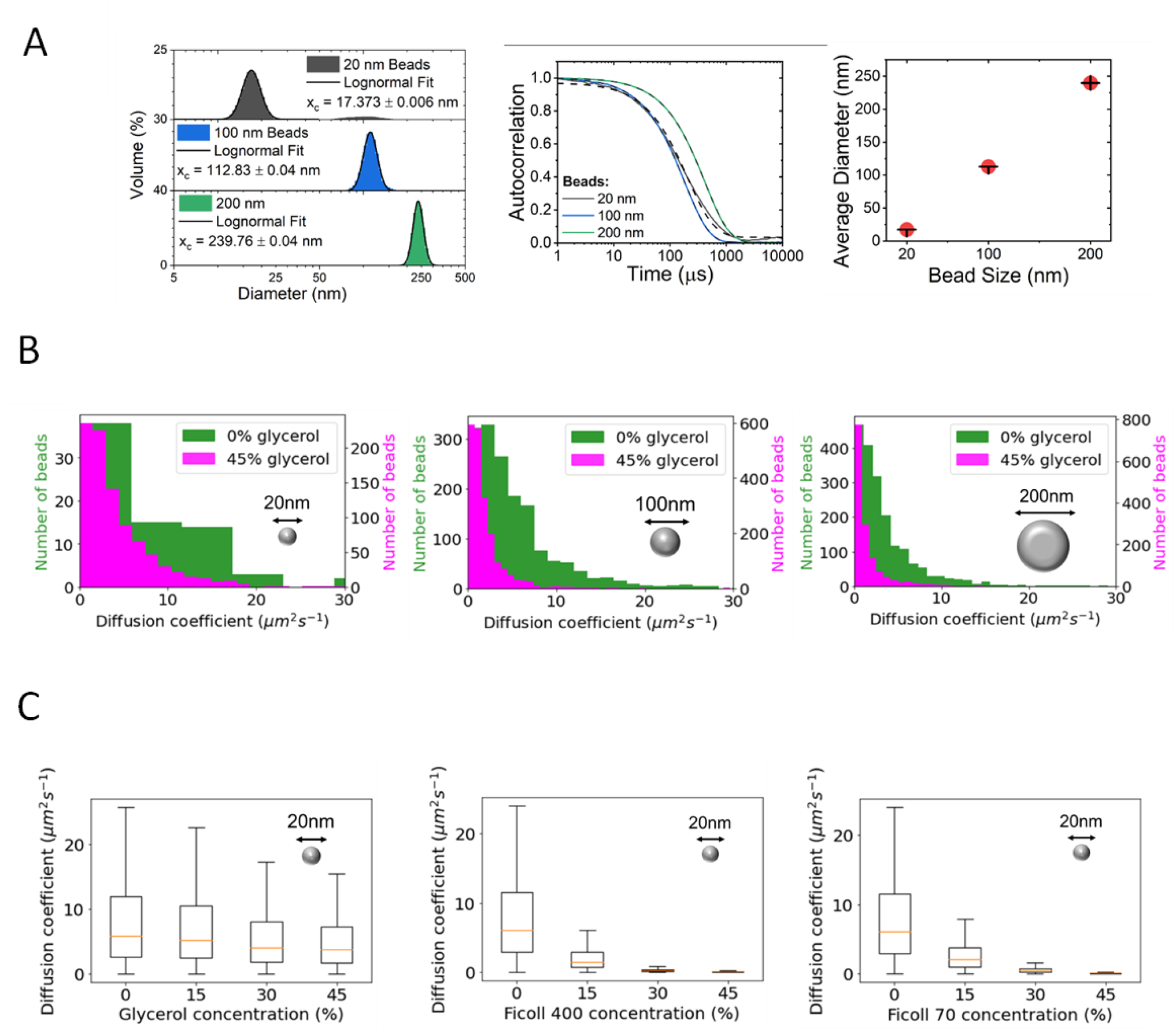
Single-particle tracking analysis of fluorescent beads *in vitro*. (A) DLS measurements for 20, 100 and 200 nm beads. (Left to right) Volume weighted distribution of particle diameter, autocorrelations for each bead type and measured bead diameters from the volume distribution fitted with lognormal distributions. (B) Distributions of diffusion coefficient of (left to right) 20 nm, 100 nm, and 200 nm beads in 0% and 45% glycerol determined through Slimfield microscopy and tracking. (C) Box plots of apparent diffusion coefficients extracted from 20 nm diameter beads diffusing *in vitro* in (left to right) glycerol, Ficoll 400, and Ficoll 70. (Log display of this data in presented in Supplementary Fig. 2.)

**Figure 3:**
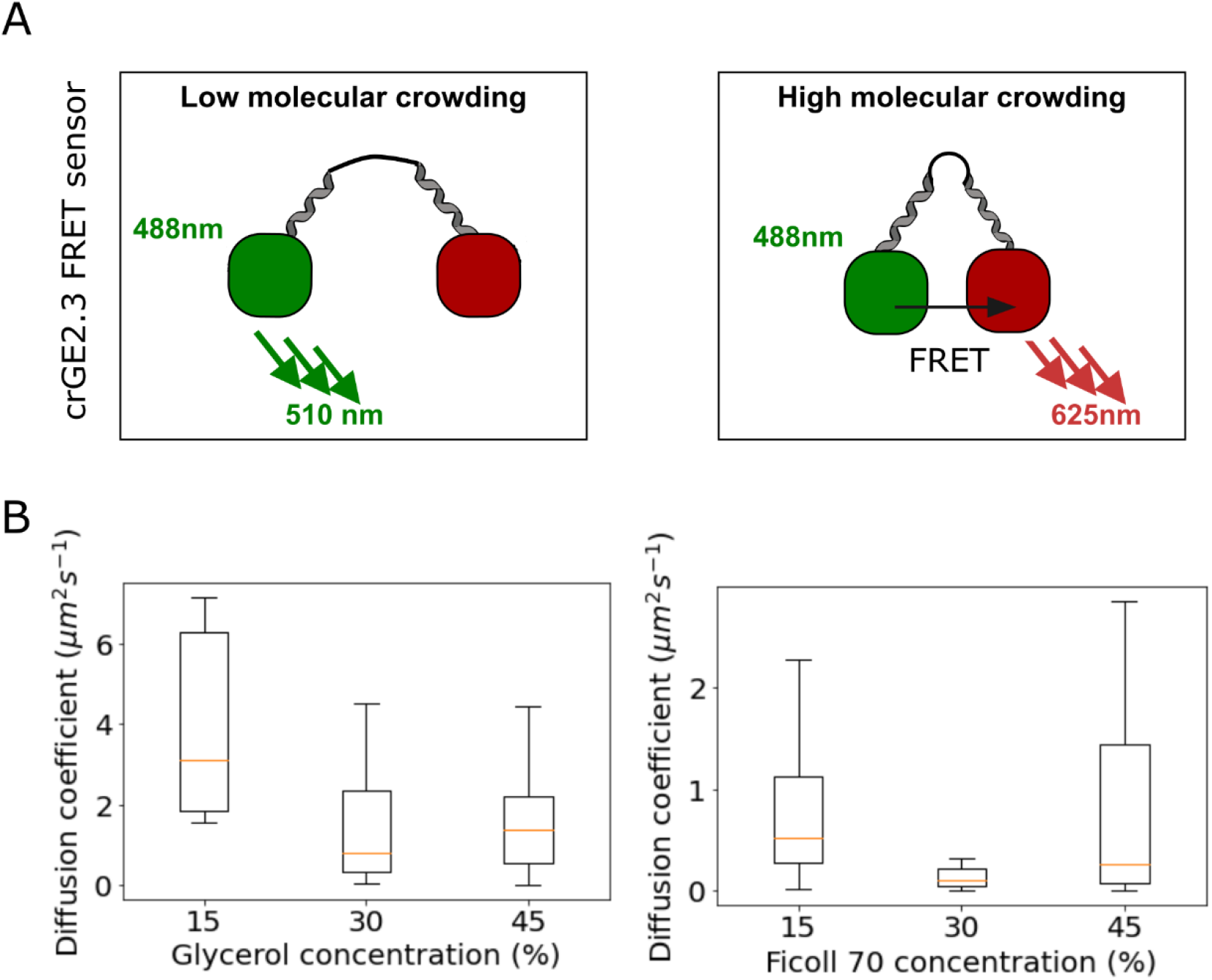
Single particle analysis of FRET sensor *in vitro*. (A) Schematic indicating the FRET based crowding sensor in different molecular crowding environments. (B) Sensor diffusion in glycerol and Ficoll 70 as determined with Slimfield imaging and trajectory analysis.

**Figure 4:**
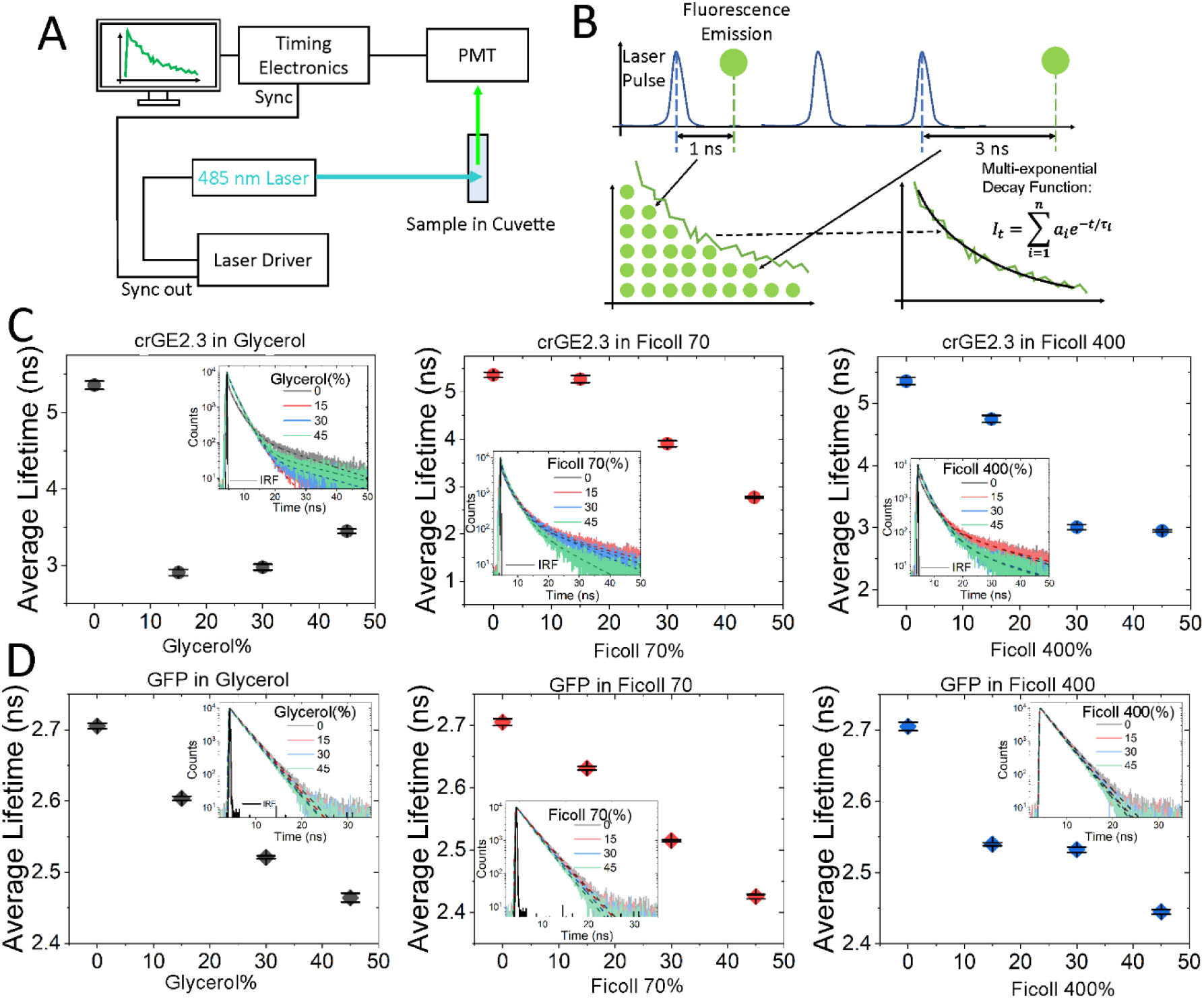
Molecular crowders change the photophysics of GFP and crGE 2.3. (A) Schematic of Time Correlated Single Photon Counting (TCSPC) building a histogram of fluorescence lifetime measurements from GFP (green circle) and fitting with a multi-exponential decay using iterative reconvolution of the instrument response function to fit the equation shown. (B) Schematic of TCSPC data acquisition. Sample in a glass cuvette is excited by a 485 nm laser from which emission is detected by a photo-multiplier tube, during this TCSPC electronics are timing how long emission takes after excitation to obtain a photon arrival time histogram representing the fluorescence decay. (C) Average lifetime of crGE 2.3 FRET crowding sensor in glycerol, Ficoll 70 and Ficoll 400. Inset: corresponding time-resolved fluorescence and instrumental response function (black). (D) Average lifetime of GFP in glycerol, Ficoll 70 and Ficoll 400. Inset: corresponding fluorescence decays and instrumental response function (black).

### 2.6 Dynamic Light Scattering (DLS) fluorescence beads measurement

DLS measurements (49) for each bead size and colour were taken with a Zetasizer mV DLS system (Malvern Instruments). This system uses the correlation of scattered laser light, of wavelength 632.8 nm, to obtain the diffusion coefficient and hydrodynamic radius through fitting a cumulant fit [1]:

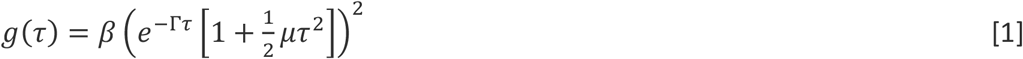

where *β* is the coherence factor, *Γ* is the decay rate, *μ* is the moment (49), and the Stokes-Einstein equation [2]:

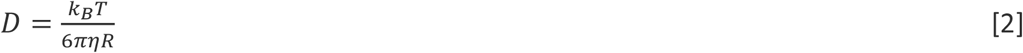

where *D* is the diffusion coefficient, *T* is absolute temperature in kelvin, *η* is dynamic viscosity and *R* is hydrodynamic radius. Each measurement consisted of 3 runs and 3 measurements were taken for each bead type, collecting back scattering at an angle of 178°. All beads were diluted (1/1k dilution from commercial beads at 2% solid) in 10mM NaPi at pH 7.4 buffer (assuming a nominal refractive index of water at 298 K of 1.33).

### 2.7 Internalising fluorescent nanobeads in *S. cerevisiae*

Wild type *S. cerevisiae* (BY4741: *MATa his3Δ1 leu2Δ0 met15Δ0 ura3Δ0*) cells were grown overnight in yeast extract peptone dextrose (YPD) liquid culture at 30°C. 5 ml of this culture was harvested and centrifuged at 3000g for three minutes, then resuspended in an Eppendorf tube in 1.4 ml of 0.2 M of lithium acetate (LiAc) solution at pH 7.5. Cells were centrifuged again at 3000g for three minutes and suspended in 100 μl of 0.2 M LiAc supplemented with 20% glycerol. A cell mix was prepared with 100 μl of cells supplemented with 100 μl of newly made LiAc master mix (4 M LiAc, 0.4 M Tris, 0.08 M EDTA, 1mg/ml salmon sperm DNA solution (UltraPure DNA salmon sperm, ThermoFisher)). 5 μl red, green and orange fluorescent beads (of indicated diameter, 20 nm, 100 nm, 200 nm) (Fluospheres, Invitrogen) were added to the mix (from a 1:1000 dilution in water). 150 μl of 58% PEG-3350 (w/v) was added to the mixture and incubated at room temperature for 30 minutes. Following this, 50 μl DMSO was added and the tubes containing cells were heat shocked at 42°C for 30 minutes. After the heat shock procedure, the cells were incubated on ice for five minutes. The cells were then washed to remove beads not internalised in the yeast. Cells were transferred into a clean glass tube, gently vortexed, transferred back in a falcon tube and centrifuged at 3,000g for 3 min, and resuspended in fresh YPD. This procedure was repeated five times so that any free beads would non-specifically adsorb to the glass tubes. Cells were also resuspended in 5 ml 1.2 M sorbitol, 10 mM sodium azide, layered on 5 ml 1.2 M sucrose, 10 mM sodium azide and centrifuged for 5 min at 3,000g as previously described (50) before resuspension in YPD and recovery growth for a minimum of 4 hours at 30°C.

### 2.8 Confocal imaging condition

Beads and yeast cells where imaged on a Commercial Zeiss confocal scanning microscope, (LSM710) equipped with a Plan-Apochromat 63× objective lens with numerical aperture of 1.4. Green fluorescence in the wavelength range 490–570 nm was collected following excitation with a 488 nm wavelength argon-ion laser, 0.1% laser power (0.037 mW at the back aperture of the objective lens). Red emissions were collected between wavelengths 570 nm and 620 nm using 561 nm wavelength excitation 0.1% laser power (0.042 mW at the back aperture of the objective lens). Z-stacks (0.37 μm slice spacing) were acquired when needed across the entire imaging volume (between the glass slide and the glass coverslip). Time-course imaging after beads internalisation was performed for 40 s at a frame rate of 1 image per second.

## 3. Results and Discussion

### 3.1 Diffusion and crowding *in vitro* depends on probe size and the specific crowding agent

To characterise the impact of viscous or crowded environments on freely diffusing molecules, we assessed the mobility of a range of fluorescent beads in various concentrations of glycerol. Two different molecular weights of Ficoll, a biochemically inert molecular crowding agent, were selected to assess the impact of crowding, Ficoll 70 and Ficoll 400. Fig. 1 shows a schematic of the bead and tunnel slide imaging method used. Three bead diameters were selected, centred on the following according to the manufacturers (but which we confirmed experimentally, see below): 200 nm, 100 nm and 20 nm, which span a full order of magnitude in diameter. Fig. 1B shows a representative DIC and fluorescence image of the beads observed with confocal microscopy. Only 100 nm and 200 nm beads were observable under confocal microscopy. 20 nm beads were too small to be resolved by DIC.

We then analysed the size distribution of the beads using DLS. We confirmed a homogeneous size distribution (Fig. 2A) as expected from commercially manufactured beads. The DLS volume distribution shows the 20, 100 and 200 nm diameter beads to have a peak diameter of 17.37 ± 0.01, 112.83 ± 0.04 and 239.76 ± 0.04 nm respectively when fit with lognormal distributions. We note that the 20 nm distribution has a secondary peak at ~100 nm which we believe is a result of a low concentration of aggregates, and this also may account for the larger than expected diameter in the larger beads. Although DLS has a greater sensitivity, complementary information may potentially be obtained from turbidity experiments in future work as a comparison.

Diffusion of 20 nm, 100 nm, and 200 nm diameter beads were measured from trajectories extracted from Slimfield microscopy micrographs for each condition. As expected, the initial diffusion coefficient of the bead has a dependence on its diameter. The well-known Stokes-Einstein relation predicts that the translational diffusion coefficient [3]:

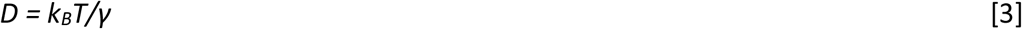

where is the Boltzmann constant, *T* the absolute temperature and *γ* the frictional drag on the diffusing object. For a perfect sphere of radius *R*, [4]

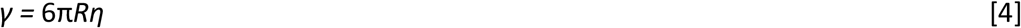

where *η* is the absolute (or ‘dynamic’) viscosity, which we subsequently just denote simply as the ‘viscosity’ (to distinguish it from the ‘kinematic viscosity’, which is *η* divided by the fluid density). In the absence of any glycerol or Ficoll, at room temperature of *T* = 298 K, the viscosity of the buffer is approximately 1 cP (or 10^-3^ Pa.s). Our diffusion measurements of the 20 nm, 100 nm, and 200 nm diameter beads in pure buffer were consistent with expectation of the Stokes-Einstein relation to within experimental error (Supplementary Fig. 1A), agreeing with a *ca*. 1/*R* dependence on *D* as predicted. We plotted 1/*R* against *D* for the buffer-only control data (see Supplementary Fig. 1C), and found that the line of best fit when constrained to pass through the origin gave a viscosity of 1.2 cP in agreement with expectations. For a further internal consistency check, we plotted mean *D* against 1/*η* in order to extract the radius of the bead from the gradient (see Supplementary Fig. 1B). We found a bead radius of 9.6 nm, in excellent agreement with the known 20 nm diameter.

The variation of viscosity with glycerol concentration has been well characterized at a macroscopic length, for example using classic Ostwald viscometry methods (51). This suggests that in the range 0-45% glycerol its viscosity compared to water might be expected to vary in the range of approximately 1 – 5 at room temperature. Similarly, although less well explored than glycerol, the viscosity for Ficoll 70 and Ficoll 400 have been characterised using ensemble viscometry methods (52). Although there are differences in how the viscosity scales with concentration and with equivalent molar mass *M* (for many dissolved polymers, including Ficoll, these can be modelled by the Mark–Houwink equation (53), where the intrinsic viscosity *η* = *KM^a^* and *K* and *a* are Mark-Houwink parameters specific to the polymer; the parameter *a* depends on molecular shape, such that solid sphere have *a* = 0 whereas random coils have a range more like 0.5–0.8 (54). Note, the Mark-Houwink equation scales the intrinsic viscosity with molecular weight, not the dynamic viscosity. Nevertheless, interested readers can find that the Mark-Houwink equation can be easily translated into a lengthscale-dependent model if they find this more instructive (55,56). These indicate a trend in both intrinsic and dynamic viscosity of glycerol < Ficoll 70 < Ficoll 400 (for example, comparing the dynamic viscosity measurements from the bulk ensembles experiments at 10% w/v indicates viscosity values of glycerol =1.3 cP, Ficoll 70 = 2.1 cP and Ficoll 400 = 4.9 cP at room temperature, in comparison to water of 1 cP), so Ficoll 70 has a viscosity which is greater than that of glycerol by a factor of ^~^1.6, whereas the equivalent factor for Ficoll 400 is closer to ^~^2.4.

Table 1 summarises our calculated diffusion coefficients for these systems. In 15% glycerol, the 20 nm diameter beads have on average higher diffusion coefficients than the 200 nm beads, as expected; the same observation applied at 45% glycerol (Fig. 2). The mean apparent diffusion coefficient, and the variance about the mean, reduced with increased concentration of either variant of Ficoll or of glycerol, though surprisingly even in high concentrations of glycerol the mobility of beads was relatively high for 20 nm diameter beads (Fig. 2B,C and Supplementary Fig. 2); the mean *D* in 45% glycerol was over half that at 0%, whereas the Stokes-Einstein relation using bulk ensemble estimates of glycerol would predict more like a 5-fold lowering of *D* over this range, with the caveat that the stochastic variation from bead to bead is clearly very high. Also, for larger beads of 100 nm and 200 nm diameter (Fig. 2B), we found that glycerol acted as a strong viscous agent strongly impairing bead mobility, suggesting that for small diffusers around the ca. 10 nm order of magnitude lengths scale, glycerol affects mobility only weakly. In Ficoll 70 and Ficoll 400, meanwhile, (Fig. 2C) even 20 nm beads are very strongly affected by low Ficoll concentrations. Surprisingly, however, the 5.7-fold increase in molecular weight between Ficoll 70 and Ficoll 400 does not result in a large difference in 20 nm diameter bead diffusion, which is consistent with their relative small reported Mark-Houwink *a* parameter of each of roughly 0.3-0.4 (52). Indeed, the two conditions give rise to a largely comparable mean and standard deviation diffusion coefficients, indicating that molecular weight itself is not a very sensitive predictor of viscosity for Ficoll variants, and crucially that viscosity and molecular crowding can be independently modulated.

**Table 1:** mean and standard deviation diffusion coefficients from *in vitro* characterisation experiments.

| Probe | Condition | Mean $D$<br>($\mu\text{m}^2\text{s}^{-1}$) | SD of $D$<br>($\mu\text{m}^2\text{s}^{-1}$) |
| --- | --- | --- | --- |
| 20 nm bead | Control | 8.42 | 8.07 |
| 20 nm bead | 15% glycerol | 7.49 | 7.25 |
| 20 nm bead | 30% glycerol | 5.74 | 5.54 |
| 20 nm bead | 45% glycerol | 5.27 | 5.05 |
| 20 nm bead | 15% Ficoll 70 | 2.87 | 2.87 |
| 20 nm bead | 30% Ficoll 70 | 0.61 | 0.64 |
| 20 nm bead | 45% Ficoll 70 | 0.12 | 0.23 |
| 20 nm bead | 15% Ficoll 400 | 2.17 | 2.20 |
| 20 nm bead | 30% Ficoll 400 | 0.35 | 0.58 |
| 20 nm bead | 45% Ficoll 400 | 0.14 | 0.43 |
| 100 nm bead | Control | 5.19 | 5.06 |
| 100 nm bead | 45% glycerol | 2.03 | 2.75 |
| 200 nm bead | Control | 3.43 | 3.69 |
| 200 nm bead | 45% glycerol | 1.61 | 2.72 |
| crGE2.3 | 15% glycerol | 5.82 | 5.95 |
| crGE2.3 | 30% glycerol | 1.43 | 1.34 |
| crGE2.3 | 45% glycerol | 1.97 | 2.51 |
| crGE2.3 | 15% Ficoll 70 | 0.85 | 0.91 |
| crGE2.3 | 30% Ficoll 70 | 0.39 | 1.65 |
| crGE2.3 | 45% Ficoll 70 | 1.54 | 2.73 |

When analysing sensor diffusion, meanwhile, we see an initial drop in diffusion in both glycerol and Ficoll 70 (Fig. 3), which then levels off as concentration increases. The mean diffusion coefficient at 15 % glycerol is approximately 2-3 fold larger than that at 45% glycerol, which is roughly as expected from the Stokes-Einstein relation using the known macro scale viscosity measurement of glycerol (51). The fluorescent protein component structures of the sensor, plus the length of the containing hinge linker region, indicate an effective hydrodynamic diameter of <10 nm. In other words, for this molecular length scale probe there is a much more sensitive scaling of diffusion coefficient with glycerol than we observed with the 20 nm diameter beads. It is worth nothing that the diffusion of single crGE2.3 molecules in 15% glycerol is lower than that of beads in equivalent conditions, likely due to higher intermolecular interactions between the organic and negatively charged protein-based sensor and surrounding macromolecules.

### 3.2 Glycerol induced viscosity and Ficoll induced crowding *in vitro* influences crGE2.3 FRET readout

We observe changes to the FRET sensor photophysics with changes to the molecular crowding environment (Fig. 4). The mGFP incorporated into crGE2.3 as a FRET pair changes its fluorescent properties when in the presence of glycerol, Ficoll 70 and Ficoll 400, to give a reduced average lifetime, suggesting that the GFP is possibly quenched in the presence of molecular crowders, potentially by an increase in energy transfer to the acceptor mScarlet-I via FRET. In this case, the observed reduction in the amplitude-weighted average lifetimes, as seen in Fig. 4C, across all crowding conditions, points towards an enhanced efficiency of energy transfer between the probes and reduced mGFP-mScarlet-I separation. Under all conditions we note that the fluorescence decays displayed three exponential components (lifetime components for each condition can be seen in Supplementary Fig. 3), which we speculate could be indicative of the sensor adopting multiple conformational states in solution. For crGE2.3 in glycerol we observed a decrease in the average lifetime of 1.9 ns, with the system exhibiting τ_av_ = 5.36 ± 0.06 ns in the absence of glycerol to 3.45 ± 0.03 ns at 45%. At high concentration of Ficoll 70, the magnitude of the decrease increased to 2.59 ns, with τ_av_ = 2.77 ± 0.02 ns observed at 45 %. A similar trend and response was then observed under Ficoll 400 conditions: the progressive addition of Ficoll 400 led to a reduction in τ_av_ to 2.95 ± 0.02 ns under high crowding conditions.

As shown in Fig.4D Free mGFP average lifetime reduces in the presence of the molecular crowders. This observation raises two points of interest. First, the observed reduction in mGFP lifetime upon addition of molecular crowders could in itself act as an indicator of molecular crowding (57–59), and second, the enhanced reduction observed from crGE2.3 upon comparison points toward an extra photophysical process, namely FRET, taking place. In other studies, the average lifetime of GFP has been seen to vary from ~1 ns to 5 ns depending on the environment (60–62) with multiple components potentially due to ground-state heterogeneity (63,64).

Here we observed an average lifetime of 2.71 ± 0.01 ns when fitted with a weighted sum of two exponential fits in the absence of molecular crowders (Fig. 4C). We then observed that the average lifetime decreased, albeit by only 0.24 ns, 0.28 ns, and 0.26 ns in 45% glycerol, Ficoll 70 and Ficoll 400, respectively. We note that the observed reduction in mGFP lifetime was ~10-fold lower than that observed in the case of the crGE2.3 FRET. This reduction is likely due to a change in refractive index when glycerol is introduced, which has been shown to decrease fluorescence lifetimes of not only GFP (64–68) but other fluorophores and fluorescent proteins such as fluorescein, CFP and YFP (69,70). Previous studies also show that Ficoll 70 and Ficoll 400 increase the refractive index of solution (52) suggesting reduction in lifetime seen for all molecular crowders used here could be due to refractive index changes.

While the observed reduction of the GFP lifetime may be associated with the refractive index change to some degree, we speculate that there may also be crowding-induced depletion interactions that lead to conformational changes within the molecule, especially on hairpin structures (71). As previously demonstrated, depletion interactions have been shown to heavily influence the structure and dynamics of simple nucleic acid switches (72), and a similar phenomenon may be occurring here on the protein hairpin-FRET based crowding sensor used in this study.

Therefore, while mGFP alone may act as a sensitive indicator of molecular crowding (57–59), these data further support enhanced FRET within the crGE2.3 system as quenchers are introduced. Further evidence to support this assertion can be seen in the emission spectra of GFP in the presence of glycerol, Ficoll 70 and Ficoll 400, as seen in Fig. 5A, where the emission of GFP is largely invariant upon addition of crowding. We note however that this could be due to direct excitation of the acceptor and therefore with higher concentrations of crowding sensor excitation spectra of the crowding sensor would further confirm that this increase in FRET efficiency.

**Figure 5:**
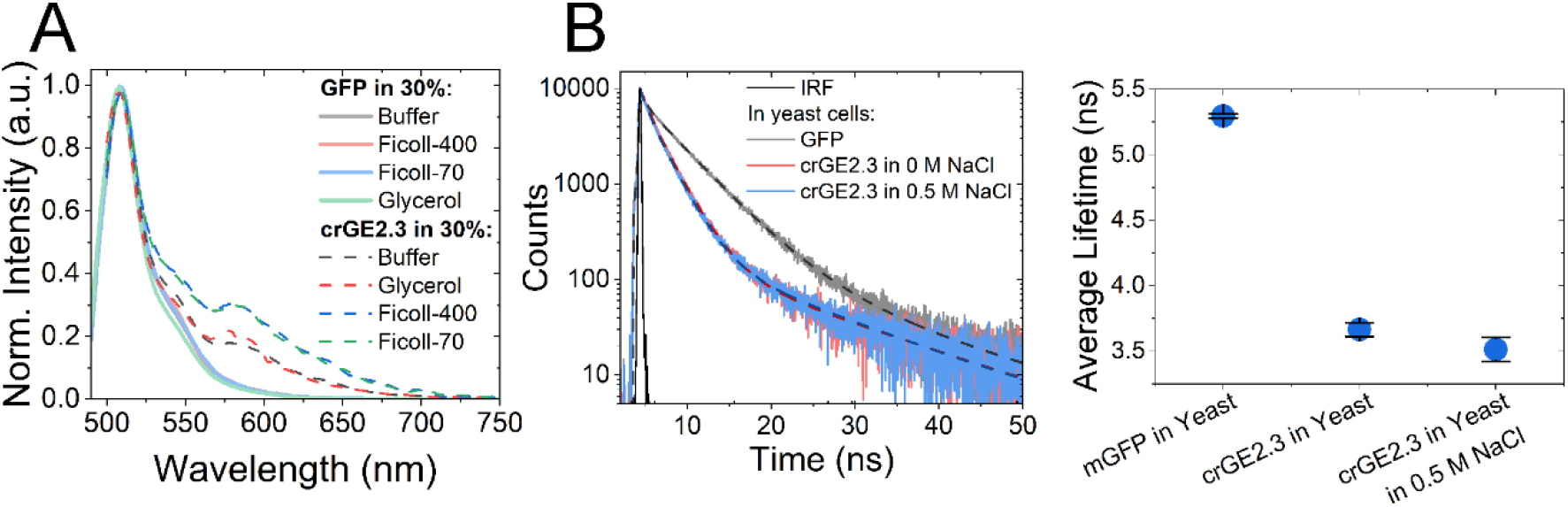
crGE2.3 spectra for FRET measurement and fluorophore lifetime characterisation in living yeast cells. (A) Emission spectra for GFP and crGE2.3 in 30% of each crowding condition. (B) Fluorescence decays for both GFP and crGE2.3 in live yeast cells and instrument response function (black) and average lifetime for crGE2.3 in live yeast cells in the presence of 0 and 0.5 M NaCl to induce stress.

However, when crGE2.3 is in the presence of Ficoll 70 and Ficoll 400 *in vitro* we observed sensitized emission at 582 nm wavelength, which is also indicative of enhanced FRET between the probes. Furthermore, the ensemble lifetime of crGE2.3 loaded live yeast cells in solution was measured, where an average lifetime of 3.67 ± 0.05 ns in the absence of cellular stress was observed (Fig. 5B). Compared to the lifetime of free GFP, this is in itself indicative of cytoplasmic crowding. Further to this, when the cells were osmotically stressed by addition of NaCl concentration at 0.5 M we observed a further shift in the lifetime to 3.51 ± 0.09 ns, pointing towards a situation whereby molecular crowding is enhanced upon NaCl stress, likely as a consequence of cellular shrinkage as the water flow out the cell (73,74).

Finally, sensor diffusion was measured *in vivo* in live yeast in unstressed conditions, yielding an average diffusion coefficient 3.8 μm^2^s^-1^, while the mean lifetime as measured with FLIM was 3.67 ns. In terms of diffusion, this makes the unstressed yeast cytoplasm approximately as diffusive as a 15% glycerol solution consistent with previous work (29,37), while the effective molecular crowding apparently lay between 30 and 45% Ficoll 70, or 15 and 30% Ficoll 400. This complexity in viscoelasticity and crowding is expected from a highly heterogeneous solution comprising macromolecules ranging in size, charge, and excluded volume, and highlights the difficulty in constructing a simplistic *in vitro* standard for comparison. We note also that the size dependence of diffusivity is dependent on the viscous agent itself, frustrating analysis. Finally, these comparisons are in terms of apparent observed behaviour only – it is impossible to say that the physicochemical environment is analogous to another on a molecular level without significant further calculation and research. It is possible only to state the behaviour on a large length- and time-scale may be similar under the conditions.

### 3.3 Method for yeast fluorescent beads up-take: *In vivo* microrehology attempt

We further attempted to characterise physical properties of the cytoplasm of yeast cells via passive microrehology particle. We focused on developing a novel method using the 20nm diameter beads that were measured *in vitro* (see section above) in our study model *S. cerevisiae*, the budding yeast. Interestingly recently other new methods with yeast (fission yeast) are emerging such as fluorescence polarization microscopy and analysis methods to access to viscosity of the vacuolar milieu (75). The following paragraph describes our method to internalised 20nm beads followed by our evaluation of the metabolic heath of the system after the procedure and limiting consequences.

We first tested cells for spontaneous or endocytic uptake when cultured in high concentration with a dense population of 100 nm and 200 nm beads. A *pep4Δ* mutant yeast strain lacking vacuolar peptidase activity was used (76,77) which might degrade beads or affect their fluorescence. Fig. 6A shows that cells fail to uptake either both 100 and 200 nm diameter beads. Z stacks through the entire volume the cells revealed that commercial beads used exhibit a tendency to non-specifically bind to both glass surfaces in the tunnel slide.

**Figure 6:**
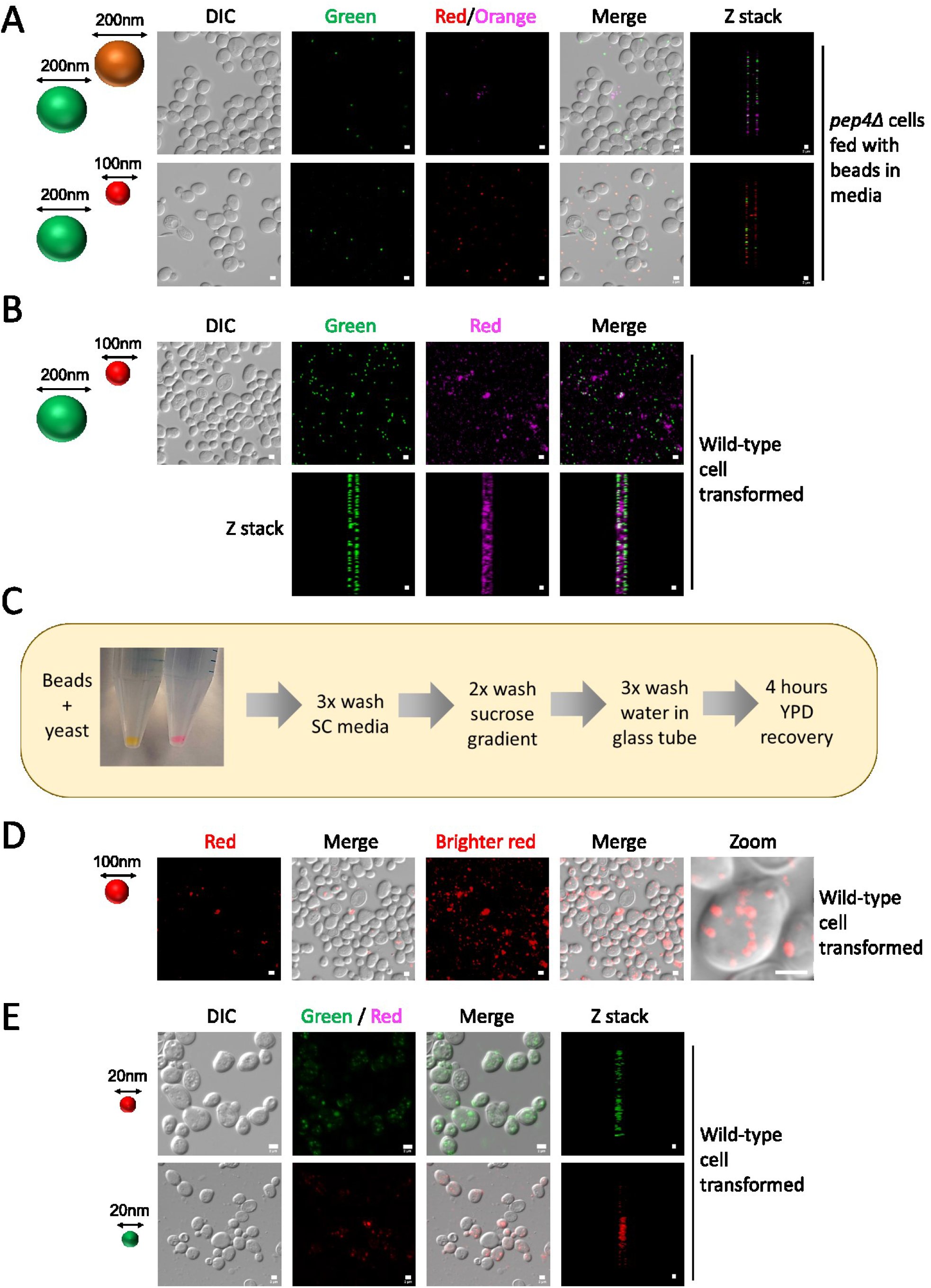
Visualisation of fluorescent bead *in vivo*, internalised in live budding yeast cells. (A) Cells lacking vacuolar peptidase activity (*pep4Δ*) were grown to mid log phase, concentrated 20-fold and incubated with labelled combinations of fluorescent beads. Cells were harvested and washed x3 in SC media prior to imaging. (B) Wild-type cells were prepared for transformation using lithium acetate (see methods) followed by heat shock. Transformed cells were then washed 3x with SC media then grown for 4-6 hours in YPD prior to imaging. (C) Flow chart for extended wash protocol following lithium acetate transformation protocol. (D) Transformation of 100 nm red fluorescent beads followed by optimized washing protocol was performed prior to confocal imaging of yeast cells. A zoomed in micrograph is included to demonstrate an example of intracellular bead organization that might imply it is associating with a structure(s) like the vacuole. E) The optimized transformation and washing protocol were performed for both 20 nm diameter red (upper) and 20 nm (lower) diameter green fluorescent beads. All scale bars: 2 μm.

We then tested transformation of 200 nm and 100 nm diameter beads into wild type cells (see Methods section). We first used electroporation which resulted in cell death of yeast decorated with fluorescent beads. More 100nm diameter beads were observed at these sites of dead cells however suggesting their uptake was more efficient (Supplementary Fig. 4). In Fig. 6B we show that 200 nm beads remain primarily adhered to the cell wall or adsorbed to the membrane, while the 100 nm beads can be seen throughout some cells’ cytoplasmic volume. The majority of beads observed stuck to glass led to set an optimised washing procedure, that included large volume of wash in glass containers were used (Fig. 6C). This optimised protocol was used to show that 100 nm diameter fluorescent beads could be transformed into yeast cells, and organised possibly through interaction with organelles like the vacuole (Fig. 6D). This protocol also was successfully used to transform 20 nm beads, both green and red labelled, into living yeast (Fig. 6E).

After successful internalisation of the beads, we measured the bead mobility using tracking, and took a confocal image every 1 s of cells containing beads. Surprisingly, this analysis showed fluorescent beads (using 20 nm diameter) remained in a static position in the cytoplasm across the imaging time scale to within experimental measurement error (Supplementary Fig. 5A). We suspect the initial period after transformation of beads, using a relatively harsh protocol, which typically allows at least 48 hours of cell recovery (78), is not conducive to physiologically relevant experimentation. Indeed, the transformation protocol resulted in morphological changes and a higher proportion of dead cells (Supplementary Fig. 5B). We grew cells for long periods (16 hours) to allow the cellular metabolism to equilibrate and cells to begin growing, but newly formed cells did not retain beads. We therefore are constrained to image cells within approximately the first 6 hours of recovery. However, this initial period is not sufficient time for transformation recovery. For example, cellular features like the vacuole, labelled with GFP-Vph1 and the mitochondria, labelled with GFP-Tom6, still exhibit gross morphology defects (Supplementary Fig. 5C). This altered physiology may be responsible for non-diffusive beads internalised in yeast. The process may alter the composition, viscosity and crowding of the cell, and the static observation may be result of a volume occupancy and physical properties of the cytoplasm altered by the transformation process. Unfortunately, cells holding beads compromise the survival and recovery process under normal growth condition preventing further investigation for cytoplasmic modulation.

## 4. Conclusions

Here, we have described a methodology for comparing the cytoplasmic viscoelasticity and molecular crowding state with *in vitro* standards through quantification of the crGE2.3 FRET-based crowding senser, and its single-molecule diffusive characteristics. We demonstrate that the diffusive landscape is related to the hydrodynamic radius of beads, as one would expect. However, and most strikingly, we found that the diffusivity of a molecular probe in glycerol was lower than that of a bead of larger extent. Also, that in the case of the glycerol, a single-molecule probe of diameter between 5 nm and 20 nm (the approximate range calculated theoretically for 2x the folded GFP radius and 2x the unfolded GFP radius) showed greater sensitivity to glycerol concentration than did the higher diameter fluorescent beads. The intermolecular interactions likely play a larger role in perturbing a small molecule probe than they do for a relatively large bead probe, the motion of which is largely dominated by thermal fluctuations of the surrounding water solvent. This result is a key consideration for microrheology – i.e., bead-based microrheology can only access bulk properties of a viscous fluid. However, the cellular cytoplasm is a complex environment which does not necessarily behave consistently across the bulk, especially at a molecular level. Through comparison with *in vitro* standards of glycerol and Ficoll we found that the *in vivo* landscape of *S. cerevisiae* is diffusively analogous to a glycerol solution, while in terms of macromolecular crowding it is more like Ficoll of varying concentrations based on molecular weights. Future work could focus on constructing a rheological standard with known molecular weight viscosity and crowding agents to faithfully reproduce the *in vivo* crowding and diffusion properties and thus tease out the specific contributions and importance of various components. However, single-bead tracking passive rheology is a useful metric of the system on a mechanical whole-cell level. To this end, we devised a method to transform 20 nm diameter beads into live budding yeast cells, previously unreported to our knowledge. We found that these beads are largely immobilised either through interactions with the cell membranes or due to the viscous environment. It is possible that the carboxylated coating may potentially have interacted with yeasts’ various subcellular organelles, and therefore further research may be beneficial to determine whether a suitable passivated bead has the same effect or not.

## 5. Author Contributions

<u>Sarah Lecinski</u>: Data acquisition, Methodology, Visualization, Writing. <u>Jack W Shepherd</u>: Data acquisition, analysis, data visualisation, Methodology, Writing. <u>Kate Bunting</u>: Data acquisition, methodology, data visualisation. <u>Lara Dresser</u>: Data acquisition, analysis, methodology, data visualisation, writing. <u>Steve Quinn</u>: Methodology, writing, funding acquisition. <u>Chris MacDonald</u>: Funding acquisition, supervision, Resources, methodology, data visualisation, Writing. <u>Mark C Leake</u>: Funding acquisition, supervision, Methodology, Resources, Writing.

## 7. Acknowledgements

Arnold J Boersma (DWI-Leibniz Institute for Interactive Materials, Aachen, Germany) and Bert Poolman (European Research Institute for the Biology of Ageing, University of Groningen, The Netherlands) for Crowding sensor crGE2.3 plasmid donation. We thank the Bioscience Technology Facility at the University of York, for support with confocal microscopy Dr Alex Payne-Dwyer and Dr Aisha Syeda for useful discussions. We thank Prof. Daniella Barillá (Department of Biology, University of York) for use of DLS instrumentation.

## 8. Data accessibility

Raw data for this article can be openly accessed from: DOI: 10.5281/zenodo.6756213 (https://zenodo.org/record/6756213).

PySTACHIO tracking software can be openly accessed via: https://github.com/ejh516/pystachio-smt/

