## supplementary methods for "Correlating viscosity and molecular crowding with fluorescent nanobeads and molecular probes: *in vitro* and *in vivo*"

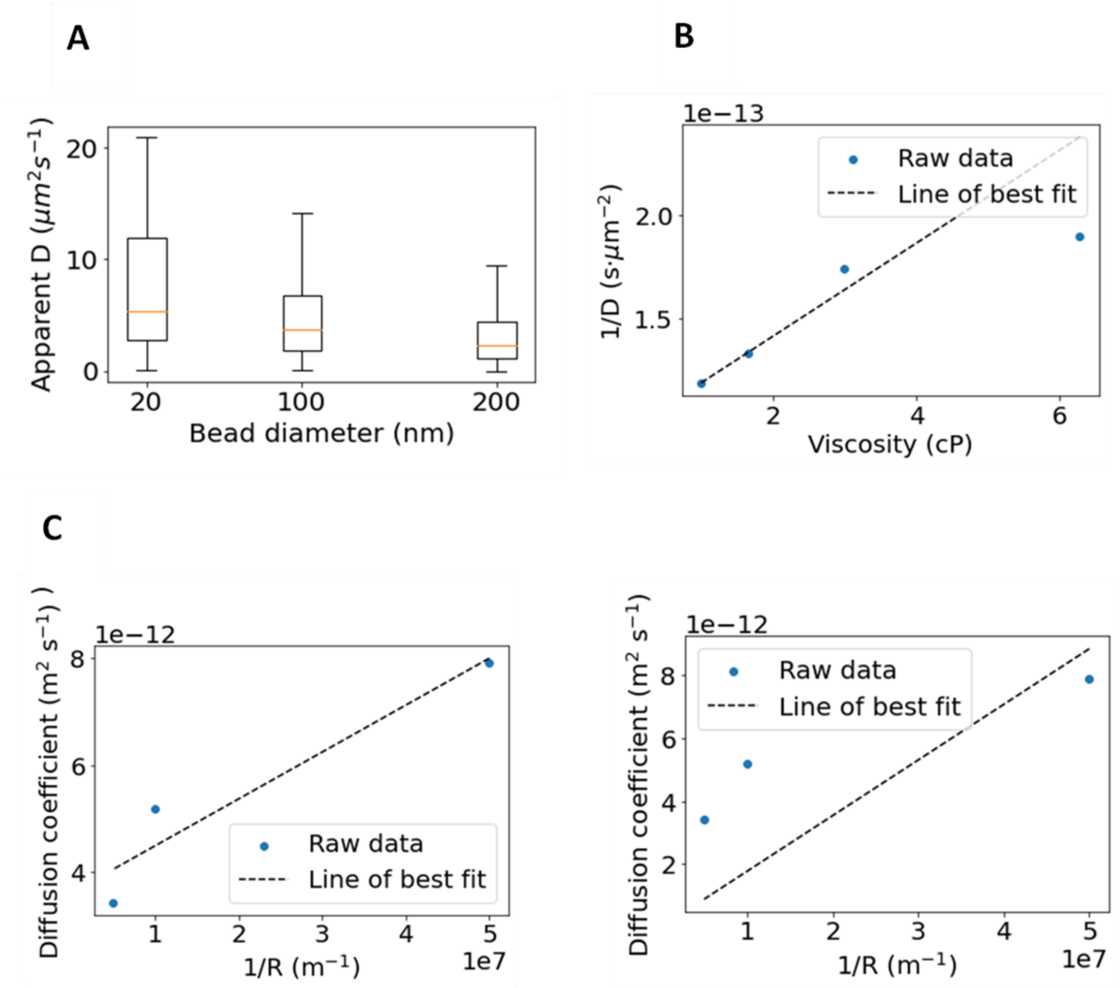


**Supplementary Figure 1:** (A) Box plot of diffusion coefficients measured for fluorescent beads of indicated diameter in the absence of glycerol or Ficoll, exhibiting the expected Stokes-Einstein relation between bead radius and diffusion coefficient. (B) Mean diffusion coefficient of 20 nm red beads in 0, 15, 30 and 45% glycerol plotted as viscosity *vs*. 1/*D* and fitted with a straight line. The gradient of the straight line fit gives an estimated radius of 9.57 nm. (C) Mean diffusion coefficient of 20, 100, and 200 nm beads plotted as 1/*R* against *D* and fitted with an unconstrained straight line. The gradient of the straight line fit gives an approximate viscosity of 2.5 cP. D) The data in C is fitted with a straight line constrained to pass through the origin. Here, the gradient gives an apparent water viscosity of approximately 1.2 cP.

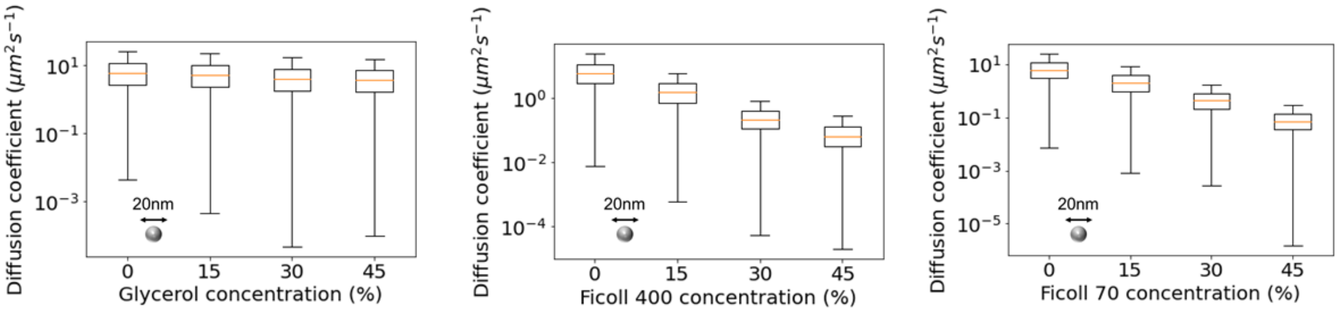


**Supplementary Figure 2 :** Box plots on a log scale of apparent diffusion coefficients extracted from 20 nm diameter beads diffusing *in vitro* in (left to right) glycerol, Ficoll 400, and Ficoll 70. (See linear plot in Fig. 2).

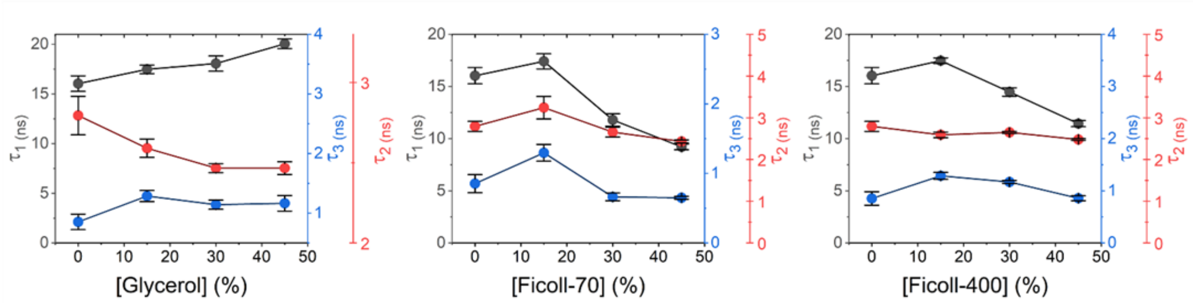


**Supplementary Figure 3:** Lifetime components of the 3-exponential fits of crGE2.3 in each condition of glycerol, Ficoll 70 and Ficoll 400.

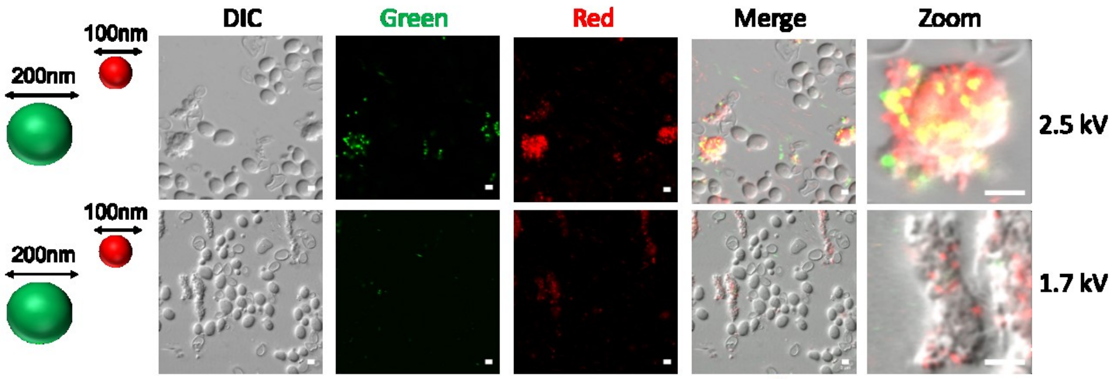


**Supplementary Figure 4:** Upper, high voltage (2.5 kV) electroporation transformation of beads results in cell death, with an association of beads with debris from particularly affected cells. Small (100 nm) diameter beads are more associated with cellular debris following electroporation than large (200 nm) diameter beads. Lower, electroporation using 1.7kV still resulted in cell death, but most cells at this less stringent transformation protocol only exhibited signs of small (100 nm) red diameter beads and not large (200 nm) diameter beads.

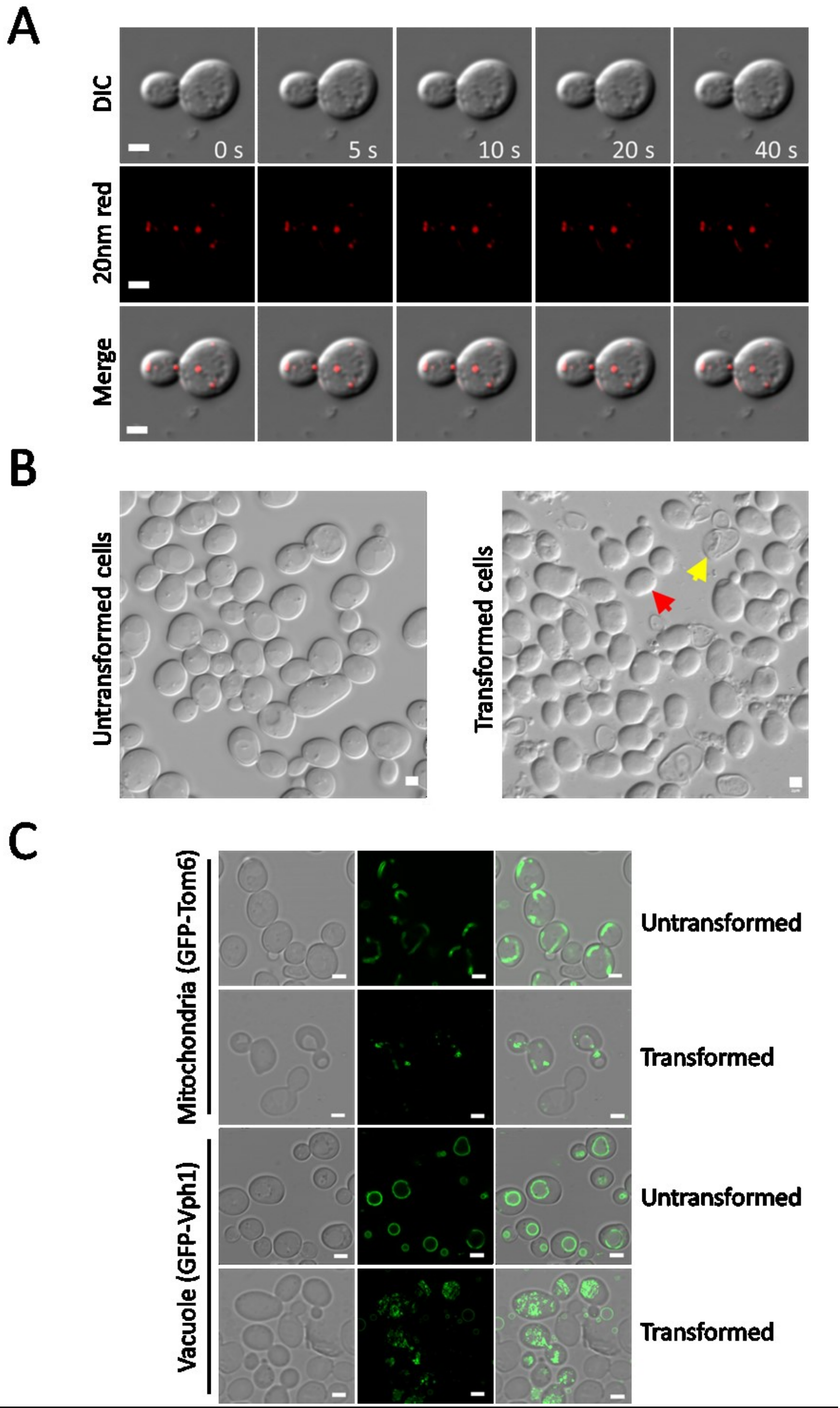


**Supplementary Figure 5: Transformed beads are immobile following transformation**

Internalised 20 nm diameter beads tracked overtime (B) Representative DIC images of control cells and yeast after transformation with the bead internalisation protocol. Yellow arrow: dead cell. Red arrow: cell stressed by internalisation protocol (C) The effect of bead internalisation protocol on the key functional organelles mitochondria and vacuole. All scales bars: 2 µm.
